# Levels of telomerase in cancer are contingent on senescence and inflammation at bulk tissue and single-cell spatial resolution

**DOI:** 10.1101/2025.05.21.655338

**Authors:** Nighat Noureen, Min Hee Kang

**Affiliations:** Cancer Center and Pediatrics, School of Medicine, Texas Tech University Health Sciences Center, Lubbock, TX, USA

**Author notes:** Correspondence: Nighat Noureen, Department of Pediatrics, School of Medicine, Texas Tech University Health Sciences Center, Lubbock, TX 79430, USA.; |.

**Keywords:** Telomerase, Senescence, inflammation, unsupervised learning, pan-cancer, single-cell, spatial transcriptomics

## Abstract

Telomerase activity plays a critical role in tumor growth and is quantified based on its level of expression. However, how these levels are associated with different pathways across various cancer types remains elusive due to the lack of a classification schema. Here, we defined an unsupervised learning metric for the quantitative measurement of telomerase activity and robustly classified the samples into low and high telomerase groups across different cancers. Using this classification system, we analyzed the data for over 9000 bulk tumors, single cells, and spatially organized tissues, and we found that telomerase high groups across the majority of cancers are strongly associated with genomic instability. On the contrary, lower group of telomerase across various cancers are significantly associated with cellular senescence, inflammation, ROS, and MAPK pathway activities. Cellular senescence, a hallmark of cellular aging, was dominant in older adults over the high telomerase levels in the majority of cancers, normal tissues, and human development phases. Our study comprehensively illustrates that lower levels of telomerase are associated with senescence-phenotype in the majority of cancers, which is strongly favorable for better survival outcomes.

## Background

Telomeres are nucleoproteins containing canonical repeats (TTAGGG) at the ends of chromosomes that play a vital role in maintaining the integrity of the linear chromosomes during cell division and prevent them from being recognized as double-strand breaks ^1^. Generally, the length of telomeres varies from 4-10 kilobases and protects the DNA erosion due to the ‘end-replication problem’ - which is defined as the lack of ability of DNA polymerase to fully replicate the terminal DNA sequences ^2^. Telomeres progressively shorten due to end replication problems by each cell division ^3^ due to oxidative stress and inflammation that cause DNA damage ^4^. Persistent shortening and dysfunction of telomeres impact cellular functions, thereby leading to senescence and cell death ^5^. On the contrary, the maintenance of telomere lengths enables bypassing senescence, thereby leading to immortality ^6^. To achieve replicative immortality, cancer cells must maintain their telomeres so that they do not shorten upon each cell division, thereby counteracting the end replication problem. The ability to enable replicative immortality by activating a telomere maintenance mechanism (TMM) is one of the hallmarks of cancer ^7^.

Approximately 85-90% of cancer cells use the ribonucleoprotein, telomerase, containing a catalytic subunit (*TERT*) and an RNA template (*TERC*) to add sequential TTAGGG telomeric repeats to the ends of chromosomes ^8–10^. The RNA subunit of telomerase, *TERC* is ubiquitously expressed across tissues ^11,12^. The catalytic subunit *TERT* is expressed in most cancers and proliferating cells. Cancer cells activate telomerase to gain immortality by acquiring *TERT* promoter mutations and rearrangements, or transcriptional dysregulation ^13,14^. *TERT* promoter mutations augment the transcriptional output of *TERT* ^15–18^ and, in certain cases, correlate with increased *TERT* expression and telomerase activity ^19,20^. Though *TERT* and *TERC* are major telomerase subunits, in many cases, *TERT* alone contributes to telomerase activity and is considered a limiting factor for it ^13–16,18,21–24^. Nonetheless, we and others have reported that *TERT* expression is not a reliable measure of telomerase activity under different scenarios ^12,25–32^. Quantification of telomerase enzymatic activities by EXTEND ^29^ revealed that each cancer type has a broad range of these activities where low and high levels across different cancer histologies are associated with patient prognosis.

Although majorly silent or extremely low in somatic cells, telomerase is active in stem cells and rapidly generating, mitotically active cells ^9,33^ and reactivated in most human cancers ^34^. Telomerase levels are heterogeneous across different cancers in multiple tissue types ^34–38^. This heterogeneity is associated with distinct tumor stages ^39,40^, molecular subtypes ^29^, and several signaling pathways. Growing evidence indicates non-canonical functions of telomerase, which affect several cellular processes, including signaling, regulation of cell survival, resistance to stress, and apoptosis ^41^. It affects cancer invasiveness and metastasis by WNT and NF-kB signaling pathways ^42,43^. The effect of telomerase on various signaling pathways is highly influenced by the quantitative levels of telomerase in different tumor samples across different tissues. Tumor cells with high telomerase activity are mitotically active with high self-renewal features and, therefore, can drive tumor growth. On contrary, tumors with low telomerase are less proliferative. Nevertheless, different levels of telomerase can trigger different signaling cascades which have a substantial impact on cellular processes like proliferation, senescence and apoptosis ^44–46^.

We and others have previously established and analyzed telomere maintenance mechanisms, *TERT* expression, and telomere lengths in cancer ^29,47–49^. However, the levels of telomerase activity have not been systematically classified and analyzed on a pan-cancer level. This study utilized an unsupervised learning approach to characterize telomerase into low and high levels across a compendium of tumor cohorts, including bulk tissue, single-cell, and spatial spectrum. We established this analysis based on our previously published telomerase activity quantification method EXTEND across cancers and normal tissues. This comprehensive integrative analysis identifies the special impact of senescence, immune inflammation, and genome instability on low and high levels of telomerase across a range of tumors in different tissues. This analysis identified that lower levels of telomerase across different tumors are significantly associated with senescence-associated phenotype, which is a suitable candidate for favorable survival outcomes.

## Results

### Unsupervised characterization of telomerase activity scores

Upon retrieving the quantitative measure of telomerase activities, computed using EXTEND, for 33 cancer histologies in The Cancer Genome Atlas (TCGA) from our previous study ^29^, we excluded tumor-associated normal cases (n=700) from this study as they have very low or negligible telomerase activities. We categorized these telomerase activity scores (EXTEND scores) into low and high classes per cancer type by implementing an unsupervised K-means clustering algorithm ^50^. This classification scheme was designed to classify the tumors robustly, using 1000 iterations over 50 random selections. The low and high telomerase groups for each cancer histology were finalized through consensus clustering of 50,000 clusters. The low telomerase activity class accounts for 57% of the TCGA cohort, while the high activity class is comprised of 43% of cases. Eleven out of the 33 cancer histologies have a significantly higher frequency of telomerase high cases (Fig. 1a, FDR < 0.05, highlighted as Group 1). This includes cancers of gastrointestinal organs (colorectal-COAD (colon adenocarcinoma), READ (rectal adenocarcinoma), STAD (stomach adenocarcinoma)), reproductive organs (uterus endometrial-UCEC, testicular carcinoma-TGCT), lung squamous carcinoma-LUSC, head and neck cancer-HNSC, thymoma-THYM, uveal melanoma-UVM and leukemia-LAML. Group 2, including kidney cancers (KICH, KIRC, and KIRP), thyroid-THCA, prostate-PRAD, and pancreatic-PAAD adenocarcinoma displayed the opposite pattern to Group1 and had 50% of cancer histologies with significant (FDR < 0.05) differences in EXTEND classes (Fig. 1a).

**Fig. 1:**
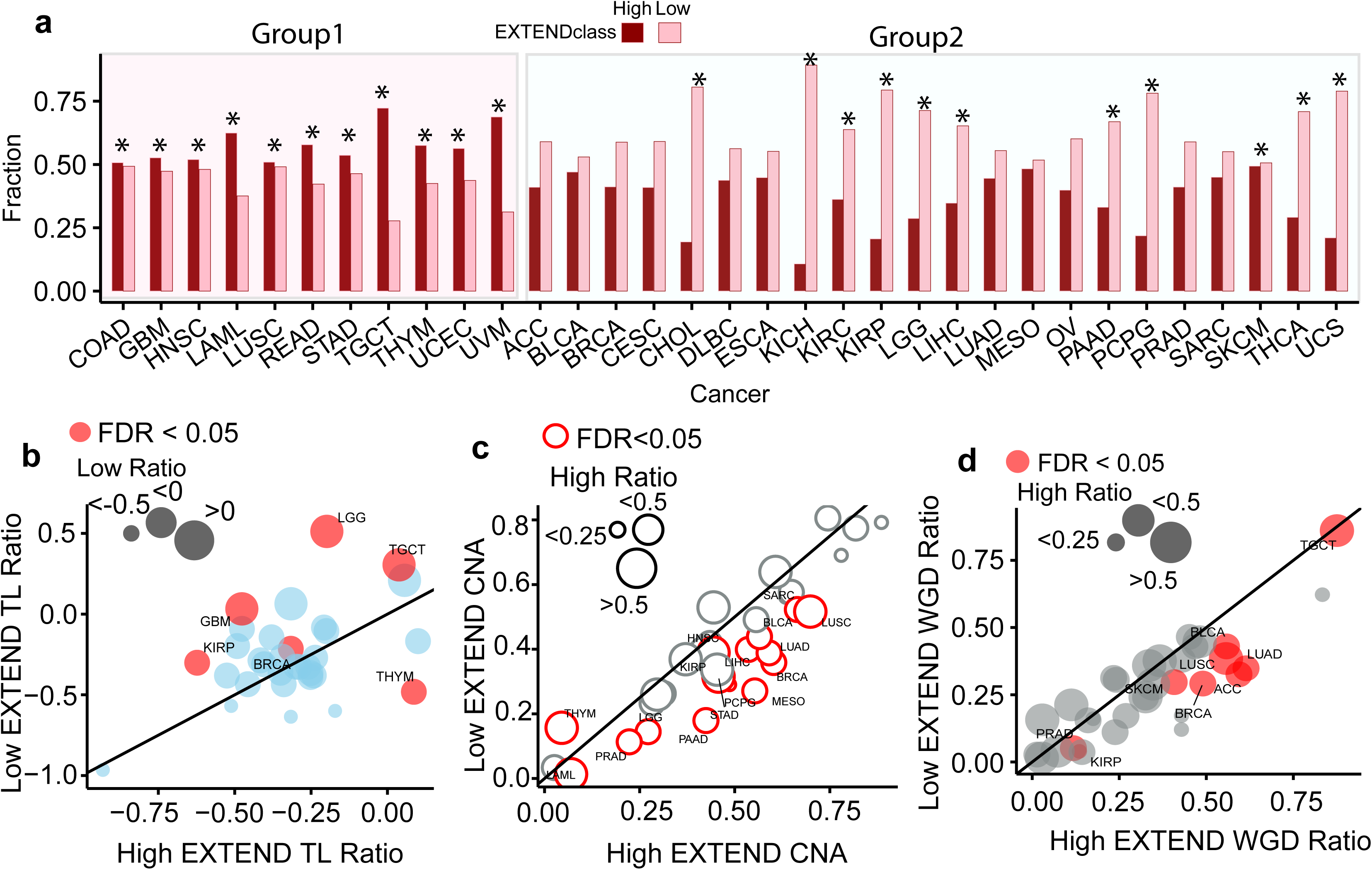
Disparities between low and high telomerase groups. **(a)** Comparison of low (pink) and high (dark red) telomerase classes in 33 TCGA cancer types. Group 1 contains cancer types with an increased fraction of high telomerase cases, while Group 2 is opposite to Group 1. Statistically significant cancer types are marked with *(FDR < 0.05). Comparisons of low and high telomerase fractions across TCGA cancer types for **(b)** Telomere length ratios **(c)** Copy number altered fractions **(d)** Whole genome doubling ratios. Cancer types are labeled for significant cases (FDR < 0.05 by Wilcox rank sum test). The data size reflects low and high EXTEND class sizes for representative cases.

### Association of clinical features with Telomerase levels

To identify characteristics associated with telomerase low and high groups, we first examined the clinical and epidemiological features of cancers in these groups. In exploring gender effects for cancers relevant in both genders, we found a significant increase in low telomerase cases in lung adenocarcinoma-LUAD (59% vs. 46%), leukemia-LAML (63% vs. 35%),melanoma-SKCM (43% vs. 33%), and head and neck cancer-HNSC (31% vs. 21%) in females representing the importance of gender for these four cancer histologies. On the contrary, higher telomerase cases were more frequent in female (39% vs 29%) in stomach adenocarcinoma-STAD (Supplementary Fig.1a). This pattern shows the association of four of the five cancers with its incidence rate being higher and survival rate being worse in males than females ^51–54^. The stomach adenocarcinoma-STAD was the only cancer type with higher cases of low telomerase activity in male patients.

The ratio of patients in the older age group (>50 years) significantly discerned high and low telomerase activity in seven out of the 33 cancers (Supplementary Fig.1b). Three cancer histologies (lung adenocarcinoma-LUAD, thymoma-THYM, and ovarian cancer-OV) showed a higher incidence (*P* < 0.05; Fisher’s exact test) of low telomerase activity in older adults than the younger patients >18 and <50 years). In contrast, glioblastoma-GBM, thyroid cancer-THCA, low-grade glioma-LGG, and uveal melanoma-UVM, showed significant (*P* < 0.05; Fisher’s exact test) increased incidence of higher telomerase activity in older adults indicating potential differences in telomerase levels among different tissue types for the younger age group ^55–57^.

We next compared telomere lengths between high and low telomerase categories across different cancer histologies. In the TCGA dataset, six out of 33 cancers showed significant differential patterns (Fig.1b, FDR < 0.05) for telomere lengths (TL). Five of the six cancers (Testicular carcinoma-TGCT, kidney cancer-KIRP, glioblastoma-GBM, low-grade glioma-LGG and breast carcinoma-BRCA) showed an inverse association of telomerase activity with TL, indicating low telomerase group bearing longer telomeres. Testis has the longest average telomere length among human tissues ^36^. Brain tumors bearing longer telomeres and low telomerase activity correspond to the ALT phenotype ^47^. The only exception to the case is thymoma-THYM, where a higher telomerase class is associated with longer telomeres.

We observed a significant association of lower disease stage (Stage I and Stage II combined) with a low telomerase class in seven out of 21 cancers (Supplementary Fig.1c, *P* < 0.05), which have sufficient sample size (n>=10). This includes adrenocortical carcinoma-ACC, liver cancer-LIHC, lung adenocarcinoma-LUAD, thyroid cancer-THCA, and kidney cancers – KIRC, KICH, and KIRP.These results corroborate with the reported literature ^39,58–60^.

We reassessed the association of patient prognosis with telomerase activity reported previously ^29^ as this study used the unsupervised classification of telomerase activity scores into low and high groups along with the exclusion of the tumor associated normals from the TCGA data. Our analysis identified 10 cancer types (Supplementary Fig.1d) exhibiting the worse overall survival for the high telomerase group including mesothelioma (MESO), thyroid cancer (THCA), pancreatic adenocarcinoma (PAAD), prostate cancer (PRAD), pheochromocytoma and paraganglioma (PCPG), kidney cancers – KIRC, KIRP and KICH, sarcoma (SARC) and adrenocortical carcinoma (ACC), whereas stomach (STAD) and thymoma (THYM) showed opposite patterns mimicking the previous analysis. The unsupervised classification excluded the lung cancer (LUAD) reported previously for the worse survival of high telomerase group.

### Genomic disparities affecting telomerase levels in cancer

To characterize genomic disparities between low and high telomerase groups, we first compared the copy number profiles, calculated based on the fraction of altered genome, and loss of heterozygosity ^61^ between these groups across different cancer histologies. Fifteen cancers had increased copy number alterations (CNAs, Fig.1c; FDR < 0.05) associated with high telomerase. Thymoma-THYM was an exception where high copy number is associated with low telomerase, a strong concordance with telomere length (Fig.1b) and survival based observations for thymoma (Supplementary Fig.1d). Lung cancers (LUAD, LUSC) were among the most prominent cases bearing significantly increased CNAs in the high telomerase class, consistent with previous literature ^47^.The assessment of loss of heterozygosity (LOH) profiles (Supplementary Fig. 2) identified thirteen cancers with high LOH associated with telomerase high class (Supplementary Fig. 2; FDR < 0.05). No association was observed for telomerase low group in LOH profile examination across pancan. Based on the strong association of high telomerase group with increased CNAs, we next reviewed the impact of whole genome doublings (WGD) on high telomerase class. We identified nine cancers with significant association between high telomerase levels and WGD, with testicular carcinoma-TGCT being the top candidate (Fig. 1d, FDR < 0.05). The other types include lung cancers (LUAD and LUSC), breast cancer (BRCA), bladder (BLCA), skin (SKCM), kidney (KIRP), prostate cancers (PRAD) and adrenocortical carcinoma (ACC) corroborating with the previous study where *TERT* expression was used as a surrogate for telomerase activity ^62^ in adrenocortical carcinoma.

Next, we compared the overall tumor mutation burden (TMB), a prominent aging-related feature ^63^ among low and high telomerase groups in the 33 cancers in TCGA. Of the 16 cancers with significant differences, 15 showed higher TMB in the high telomerase group (Fig. 2a, FDR < 0.05) while thymoma-THYM had higher TMB in the low telomerase group. These results are in concordance with the survival pattern (Supplementary Fig.1d) where high telomerase in THYM is associated with better survial than low telomerase group. To investigate the association of low and high telomerase groups with frequently mutated genes across Pancan, we calculated the percentage of recurrent gene mutations (n = 16, Fig. 2b) found in two or more cancers (n=17). Most recurrent gene mutations showed a lower mutation rate in the low telomerase group, consistent with the overall TMB of low telomerase across Pancan. The most frequently mutated gene was *TP53,* which showed recurrent mutations in nine cancers for the high telomerase group and in only three cancer types (THYM, LGG, and HNSC) for the low telomerase groups. Intriguingly, *KRAS*, *PKHD1L1*, *RB1,* and *TTN* genes showed recurrent patterns only in the high telomerase groups, while *PTEN* and *ATRX* genes showed recurrence in low telomerase groups, thereby representing ALT phenotypes across different cancer types. The non-recurrent and cancer-specific genes (Fig. 2c) include 28 genes across 11 cancers where higher mutation percentage mostly correlates with high telomerase activity with exceptions, including *PPP2R1A* in UCS, *HRAS,* and *GTF2I* in THYM and *SMAD4* in PAAD. The *CIC* gene showed a significantly higher association with the telomerase high group in LGG, hence highlighting *TERT* high expressing cases. This significantly maps to the fraction of *TERT* promoter mutations, which are higher in the high telomerase group in LGG and THCA (Fig. 2d). On the contrary, *ATRX* gene mutations were higher in the low telomerase group in LGG, SARC, and GBM (Fig. 2e) marking the ALT phenotype.

**Fig. 2:**
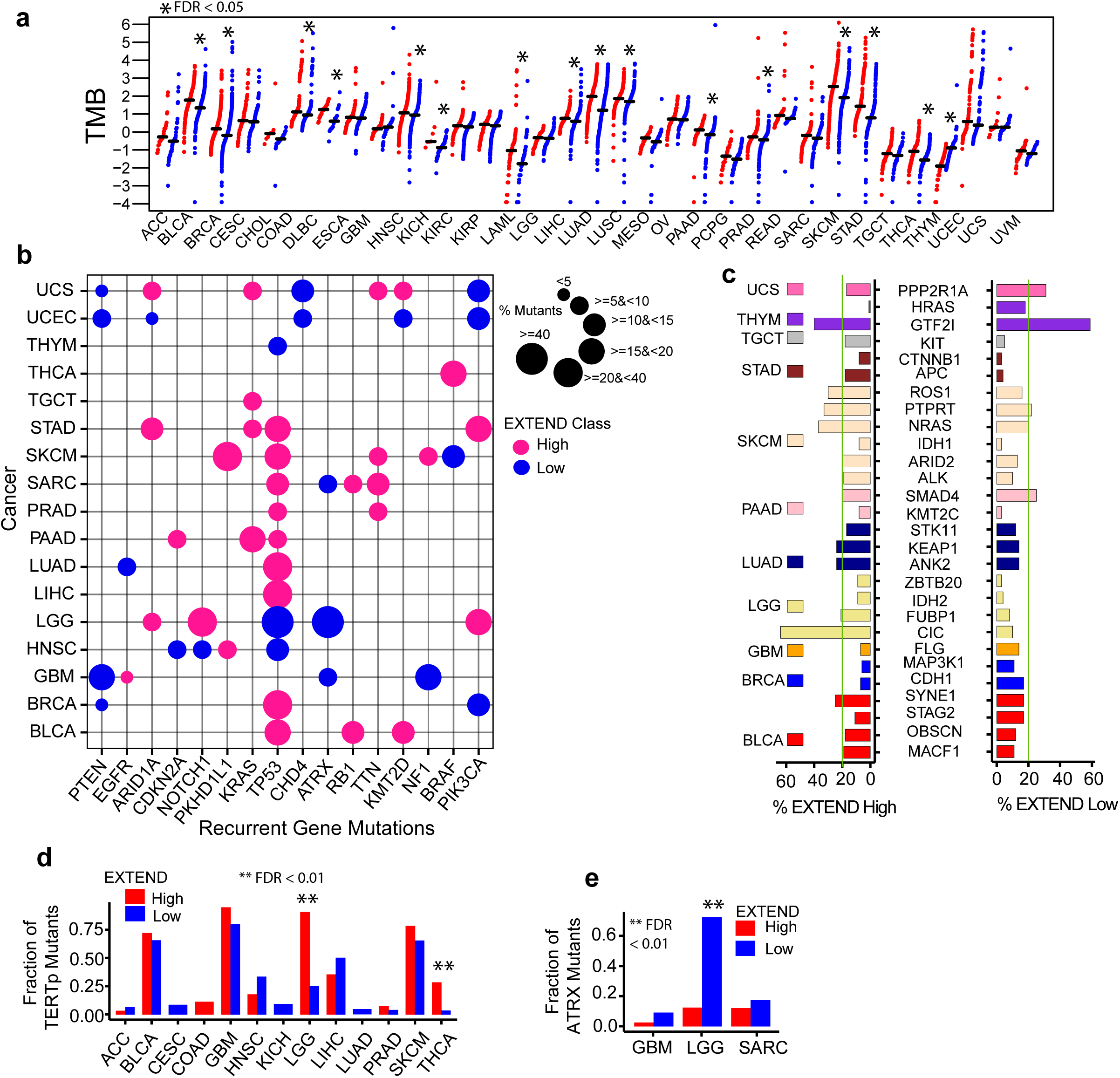
Mutational differences among High and Low Telomerase Groups. **(a)** Tumor mutation burden across 33 cancer types. Red reflects high, and blue reflects low telomerase (EXTEND) class samples. Significant ones (FDR < 0.05, student’s t-test Pvalue corrected for fdr) are marked with an asterisk **(b)** Percentage of recurrent genes (n=16) mutations across 17 cancer types in high (pink color) and low (blue color) telomerase groups. **(c)** Cancer-specific gene mutations (n =28) percentage in 11 cancer types across low and high telomerase groups. Differential patterns of **(d)** TERT gene promoter mutations across 13 cancer types **(e)** ATRX gene mutations across three cancer histologies, in low and high telomerase groups. Significant ones are shown with an asterisk (FDR < 0.01).

To understand the mechanism of increased TMB and CNAs in telomerase-high tumors, we next examined the gene set pathway enrichments for high and low telomerase groups. In pathways from Molecular Signatures Database ^64,65^, the telomerase-high group was associated with stemness and proliferation-related pathways across TCGA Pancan. On the contrary, the telomerase low group was mainly linked with muscle-associated pathways, a feature of differentiated skeletal muscles. The most intriguing top hit was MAPK targets, which were up-regulated in the low telomerase group (Supplementary Tables 1 & 2).

We also investigated the association of telomerase groups with gene fusions ^66^ that play a critical role in oncogenesis. Based on our initial observations, median fusions in TCGA associated with telomerase levels were very low (Supplementary Fig. 3a), with few exceptions. This includes seven cancer types (ACC, BRCA, KIRC, LUAD, OV, PRAD, and READ) where the high telomerase group showed a significant association with increased fusion frequencies (Supplementary Fig. 3a; *P* < 0.05). The recurrent gene fusion analysis identified 18 cancer types with one or two gene pairs with recurrent fusions, while four cancer histologies (esophagus-ESCA, glioblastoma-GBM, leukemia-LAML, and prostate cancer-PRAD) with >2 gene pairs, leukemia (LAML) being the most prominent one with eight vs. five gene-fusion pairs in high and low telomerase levels, respectively (Supplementary Fig. 3b). Recurrent gene fusion pairs were selected if >1% of samples per cancer type contained the fusion. Of these gene-fusion pairs, 22 showed specificity towards the high telomerase group while 15 were low telomerase-specific (Fig. 3a). We identified nine gene pairs with differential patterns in low and high telomerase classes (Fig. 3a). Among them, three cases were associated with the high telomerase class and six with low telomerase class. The higher frequency of recurrent gene fusions in the high telomerase class indicated its association with higher levels of telomerase. Among them, the significant ones (*P* < 0.05) include IRF2BP2-SP2 (PCPG), MECOM-LRRC31 (ovarian cancer-OV), and NKD1-NOD2 (low-grade glioma-LGG) ^67^ for high telomerase associated cases. The significant cases of low telomerase-associated recurrent gene fusion pairs include PML-RARA, KMT2A-MLLT10, and ABR-YWHAE in leukemia (LAML) ^68,69^, CCDC6-RET ^70,71^ in thyroid carcinoma (THCA), SLC45A3-ERG ^72^ in prostate adenocarcinoma (PRAD) and FGFR3-TACC3 in lung squamous carcinoma (LUSC). For fusions associated with drug targets^66^, we identified 27 drug targets (both off-label and on-label) associated with 29 cancers in >1% of the samples (Fig. 3b). Of these, 31 were high telomerase-specific, and 40 were low telomerase-associated across different cancers (marked with green X, Fig. 3b). The *TMPRSS2* and *MET* gene fusions were significantly (*P* < 0.05) associated with high telomerase groups in prostate cancer (PRAD) and kidney cancer (KIRP), respectively. On the contrary, *ROS*, *RET*, *PML-RARA,* and *FGFR3* gene fusions were significantly high in lung adenocarcinoma (LUAD), thyroid cancer (THCA), leukemia (LAML), and lung squamous carcinoma (LUSC), respectively for low telomerase cases.

**Fig. 3:**
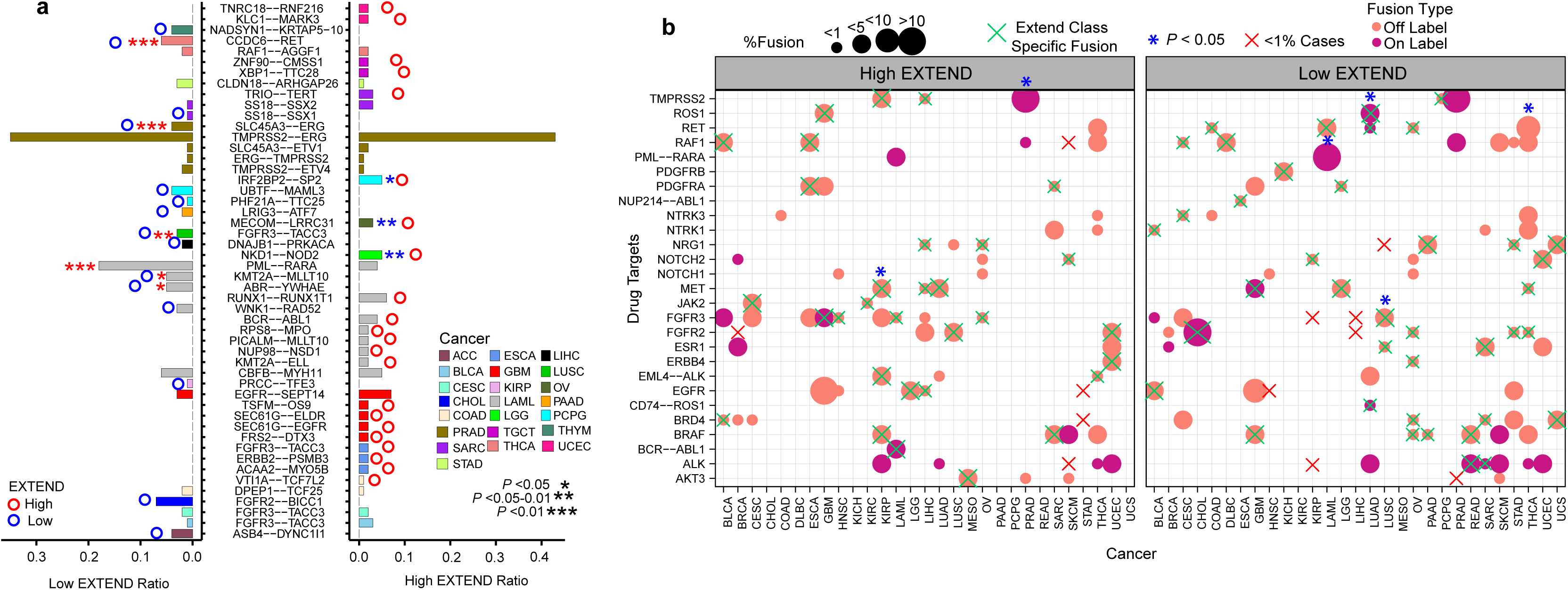
Variations in gene fusions in Low and High Telomerase Groups. **(a)** Recurrent gene fusion fractions across TCGA pan-cancer for low and high telomerase groups. Fusions with more than 1 percent of cases per cancer are included in the analysis. Barplots show the ratio of fusions in each EXTEND class. Asterisk marks significance level (P < 0.05), blue (low EXTEND), and red (high EXTEND) circles on top of some bars show EXTEND class-specific gene fusions. **(b)** Association of druggable gene fusion targets with low and high telomerase groups. Targets with greater than 1 percent cases per cancer type are included in the analysis. *(blue) shows significantly differential cases between low and high telomerase (Fisher’s exact test P < 0.05). EXTEND class-specific fusions are marked with a cross (X in green) and in red if the sample size is less than 1 percent for a particular telomerase group. Data point size reflects % fusion cases. The color code of data points represents fusion type, including both off-label and on-label fusions.

### Association of Senescence with Telomerase Levels in Cancer

Based on increasing evidence, the role of senescence in cancer is controversial. Its tumor-promoting characteristics promote it as an emerging hallmark of cancer ^73–76^. Though the senescence phenotype possesses low proliferative potential ^77^, they can escape senescence arrest and undergo continuous division ^78^ by adapting the telomere maintenance mechanism, which in most cancers is attained by reactivating telomerase ^79^. Since senescence and telomerase in cancer are tightly interconnected, we examined the association of cancer senescence and levels of telomerase across different cancer types.

We computed senescence scores (details in methods) for 33 TCGA cancer types using a recently published senescence signature ^80^. This signature was calculated from an expression signature of 125 genes, primarily consisting of senescence-associated secretory phenotype (SASP) factors and intracellular and transmembrane proteins. We found senescence being the marker of non-proliferative cells was significantly (FDR<0.05) associated with the low telomerase groups in 26 cancers(Fig. 4a, FDR<0.05). We also observed a significant negative correlation of telomerase with senescence in pediatric neuroblastoma ^81^ (P=2.3e-07, Supplementary Fig. 4a) and cancer cell lines data from CCLE (Fig. 4b, *P* < 2.2e -16) thereby arbitrating high senescence in low telomerase groups across a variety of cancer or non-cancer cell lines (Supplementary Fig. 4b), with fibroblasts being the top senescence candidates with low telomerase activity and hematopoietic lines having the lowest senescence and higher telomerase scores due to their self-renewal potential ^82^.

**Fig. 4:**
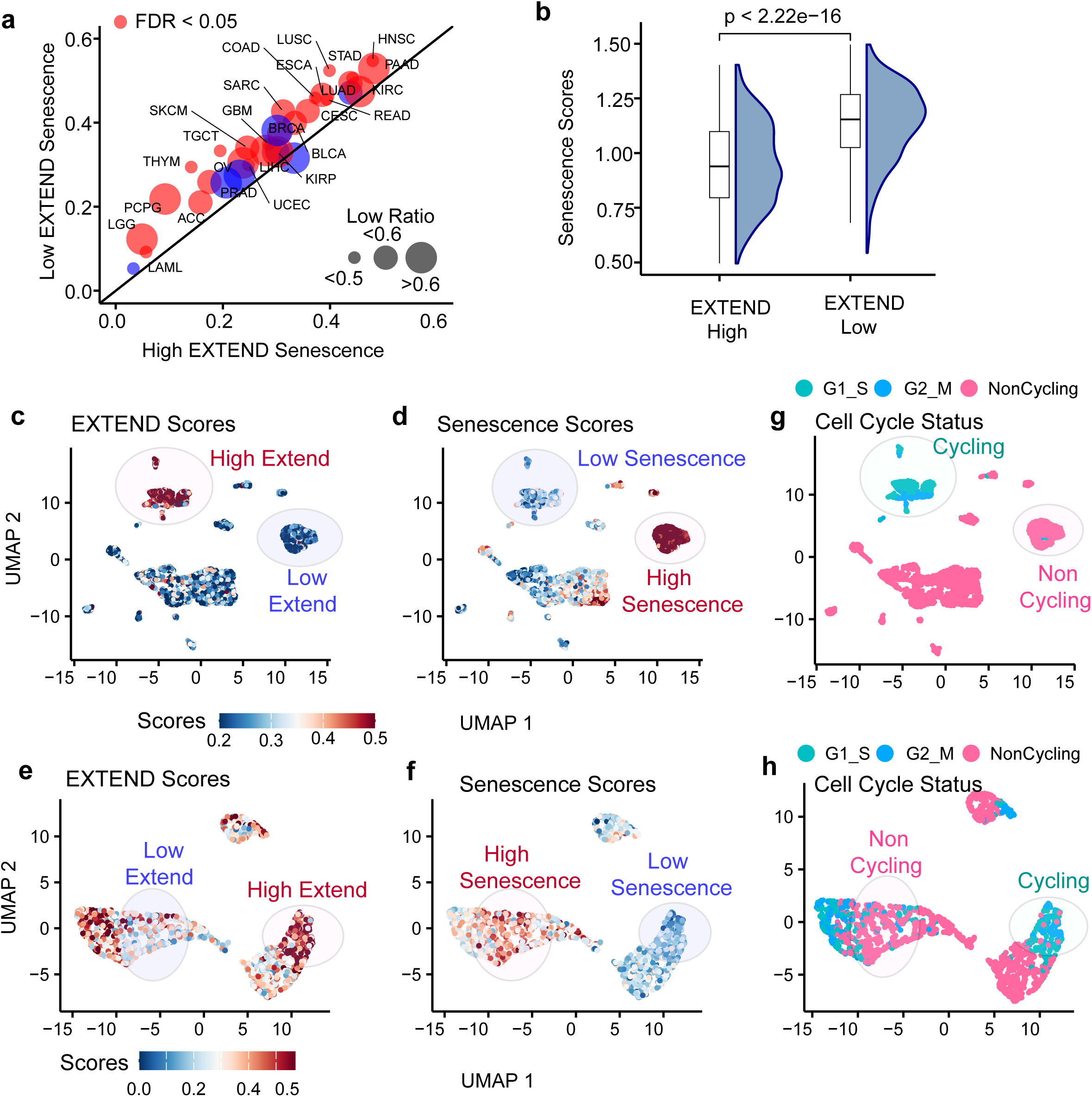
Senescence scores distribution in low and high telomerase groups. **(a)** Differential patterns of senescence scores across low and high telomerase groups for TCGA pan-cancer data, size represents low EXTEND ratio, color (red) represents significance (FDR < 0.05). **(b)** Senescence scores differences among low and high EXTEND classes using Student’s T-test for cancer cell lines data (CCLE). UMAPs representing the alignment of high and low telomerase (EXTEND) scores/groups with low and high senescence scores in **(c-d)** Single-cell GBM data (Neftel et al., 2019) and **(e-f)** Single-cell HNSC data (Puram et al., 2017). Cell cycle groups aligned for UMAPs in c-d for **(g)** single cell GBM data and in e-f for **(h)** single cell HNSC data.

We next sought to investigate if the tumor and tissue level patterns reflected the tumor cell behavior. We calculated senescence scores using single-cell RNAseq data from glioblastoma ^83^ and head and neck cancer ^84^. Using the EXTEND scores for the single cell data sets, we again observed a strong association of senescence with the low telomerase group (Figs. 4c-d and Figs. 4e-f and Supplementary Fig. 5a and Supplementary 5b; *P* < 2.2e-16) signifying innate reverse correlation between two pathways in cancer cells. Distribution of senescence scores across cell cycle phases for single-cell data highlighted significantly higher activity in noncycling cells in both GBM (Fig. 4g; Supplementary 5c, *P* < 2.2e-16) and HNSC (Fig. 4h; Supplementary 5d, *P* < 2.2e-16) thereby confirming the reverse correlation pattern of senescence with active cell cycle at cellular resolution.

To mark the reverse correlated properties of telomerase and senescence in different sections of the cancerous tissue, we investigated their spatial settings in two spatial transcriptomic data sets from lung ^85^ and breast ^86^ tissues. We calculated telomerase, senescence, and cell cycle scores for both the spatial transcriptomics datasets. We then compared the enrichment of different sections of these tissues for telomerase, senescence, and cell cycle scores. The enrichment patterns truly reflect the opposite patterns of telomerase and senescence in various sections (Fig. 5a-5b, 5c and 5d) of both tissues, mimicking the reverse correlation observed at bulk and single-cell RNAseq levels. The cell cycle activity, like the telomerase activity, showed the anticorrelated patterns to the senescence in spatial sections of the lung (Fig. 5e) and breast tissue (Fig. 5f).

**Fig. 5:**
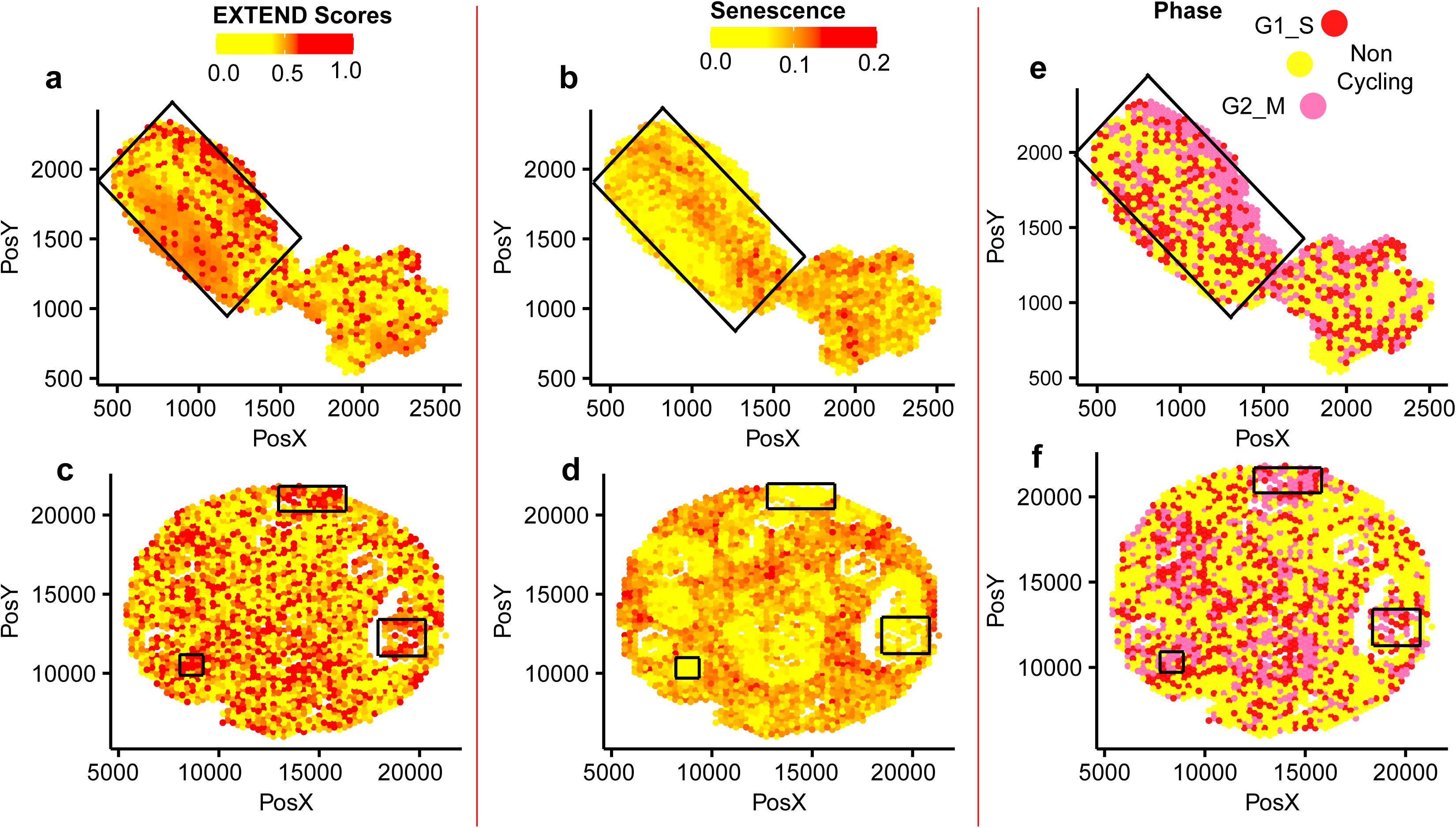
Telomerase comparisons in spatial settings. Telomerase comparisons in spatial settings. EXTEND versus senescence scores enrichment across two spatial transcriptomics data sets **(a-b)** Lung cancer **(c-d)** Breast Cancer. Cell cycle phases correlated enrichment mapping to senescence and telomerase activity scores for spatial **(e)** Lung cancer data set **(f)** Breast cancer data set. The highlighted black boxes for all cases reflect enriched patterns for high telomerase, low senescence, and high cycling activity and vice versa from a to f.

These observations reflect that high telomerase activity or its reactivation attenuates senescence levels in tumors at both tissue and cellular levels. Though well-known previously, tumor cells must bypass senescence to maintain telomere lengths by reactivating telomerase ^87^. Nevertheless, it has not been quantitively delineated at a pan-cancer level, especially in spatial and single-cell settings.

### Senescence is negatively associated with age in the context of telomerase

To compare senescence with age in the context of telomerase activity, we took the cancer types with sufficient sample size (n>50) that emcompass 21 cancers from TCGA data where we could distinguish older adults (OA). Comparison of senescence scores between high and low telomerase in OA patients demarcate significant differential patterns (Fig. 6a, *P* < 0.001). These patterns mimic the senescence bypass mechanism and reactivation of telomerase in high versus low telomerase groups across 21 cancers (Fig. 6a) due to higher levels of *TERT* expression resulting from promoter mutation or methylation^88^.

**Fig. 6:**
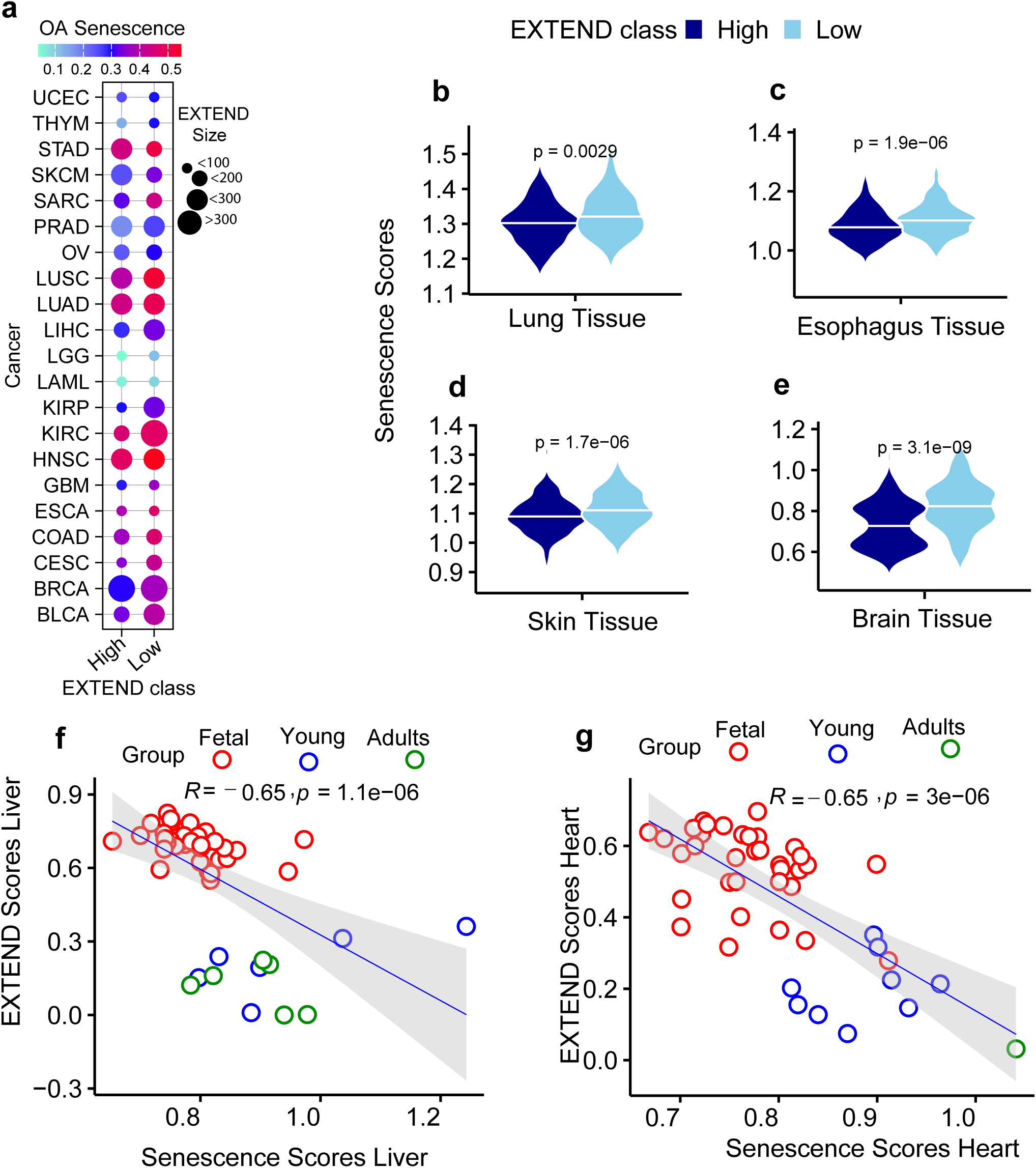
Senescence and telomerase disparities based on age groups. Significant differential patterns of Senescence in older adults for low and high telomerase class **(a)** in TCGA cancer types, the size of data represents low and high telomerase class size, and the color panel shows senescence for OA in each cancer type. Cancer types with non-significant levels are not included. In GTEX tissues **(b)** Lung **(c)** Esophagus **(d)** Skin **(e)** Brain, *P* values are calculated using T-test. GTEX-only tissues taken with a sample size > 50 in both cases and are present in TCGA data. Correlation of Senescence and EXTEND scores in human development for **(f)** Liver **(g)** Heart, tissues. Color represents age groups (red=fetal, blue= young, and green=older adults).

Based on the data from 21 TCGA cancers, we investigated senescence levels for the same tissues in normal tissue data obtained from Genotype Tissue Expression (GTEx portal). We identified four tissues with sufficient sample size for OA cases, including lung, esophagus, skin, and brain tissues. Normal tissues mimic the senescence mechanism as shown in tumor tissues with significant differential patterns in these four tissue types (Fig. 6b-e, T-test *P* < 0.001).

We next examined the association of telomerase activity with senescence in human liver and heart tissues for their diverse self-renewal capacities during embryonic development ^89^. We computed senescence scores for heart and liver samples and compared them to telomerase activities. Due to the low sample size at each developmental phase, we did not divide them into subgroups. We measured the association of telomerase and senescence, which captured the negative correlation between two factors in each tissue (Fig. 6f-6g, *R* = -0.65 and *P* < 0.0001). The significant high telomerase and low senescence in fetal samples in both heart and liver tissue indicate active cell cycle and stemness behavior. On the contrary, this pattern is opposite in younger and older adults, where an increase in senescence scores is observed with a decrease in telomerase activity, thereby confirming the loss of proliferative mechanism in later age groups ^90^. Liver tissue for younger and older adults showed a mixed pattern because of active self-renewal and active telomerase at those stages ^91,92^.

### Immune inflammation in cancer is associated with low levels of telomerase

Telomere attrition is associated with an active cell cycle hence an active telomerase in cancer tissues ^47^. As the active telomerase is associated with active proliferation and stemness across pan-cancer ^29^, up-regulation of telomerase facilitates immune response and prevents immune cell senescence ^93^. Therefore, we investigated how telomerase levels would modulate immune response or vice versa.

We obtained immune subtypes of TCGA tumors from a previous pan-cancer analysis ^94^. These subtypes were calculated based on various measures, including immune signatures, macrophage/lymphocyte levels, proliferation rates, intra-tumoral heterogeneity, and other immune features. We found that low levels of telomerase in 18 cancer types were significantly (Fig. 7a, Fisher exact Test FDR < 0.05) associated with the C3 immune subtype. This suggests that C3, being the low proliferative class of tumors, is significantly associated with low telomerase activity. Fewer exceptions were observed for some cancers, with a certain fraction of samples mapping to high proliferative immune classes C1 and C2. Testicular carcinoma (TGCT) was unique, with high and low classes of telomerase significantly mapped to high proliferative class, indicating an overall active cell cycle in the testis. The lymphocyte-depleted C4 immunologic class distinctly associated sarcomas (SARC) and adrenocortical carcinomas (ACC) with high telomerase along with three other cancer types. None other than some samples from the kidney (KICH) were mapped to the immunologically quiet class, highlighting indolent cases. Very few samples of low telomerase from liver, lung, mesothelioma, pancreas, stomach, and breast cancers mapped to the TGF-B-dominant C6 immunologic class.

**Fig. 7:**
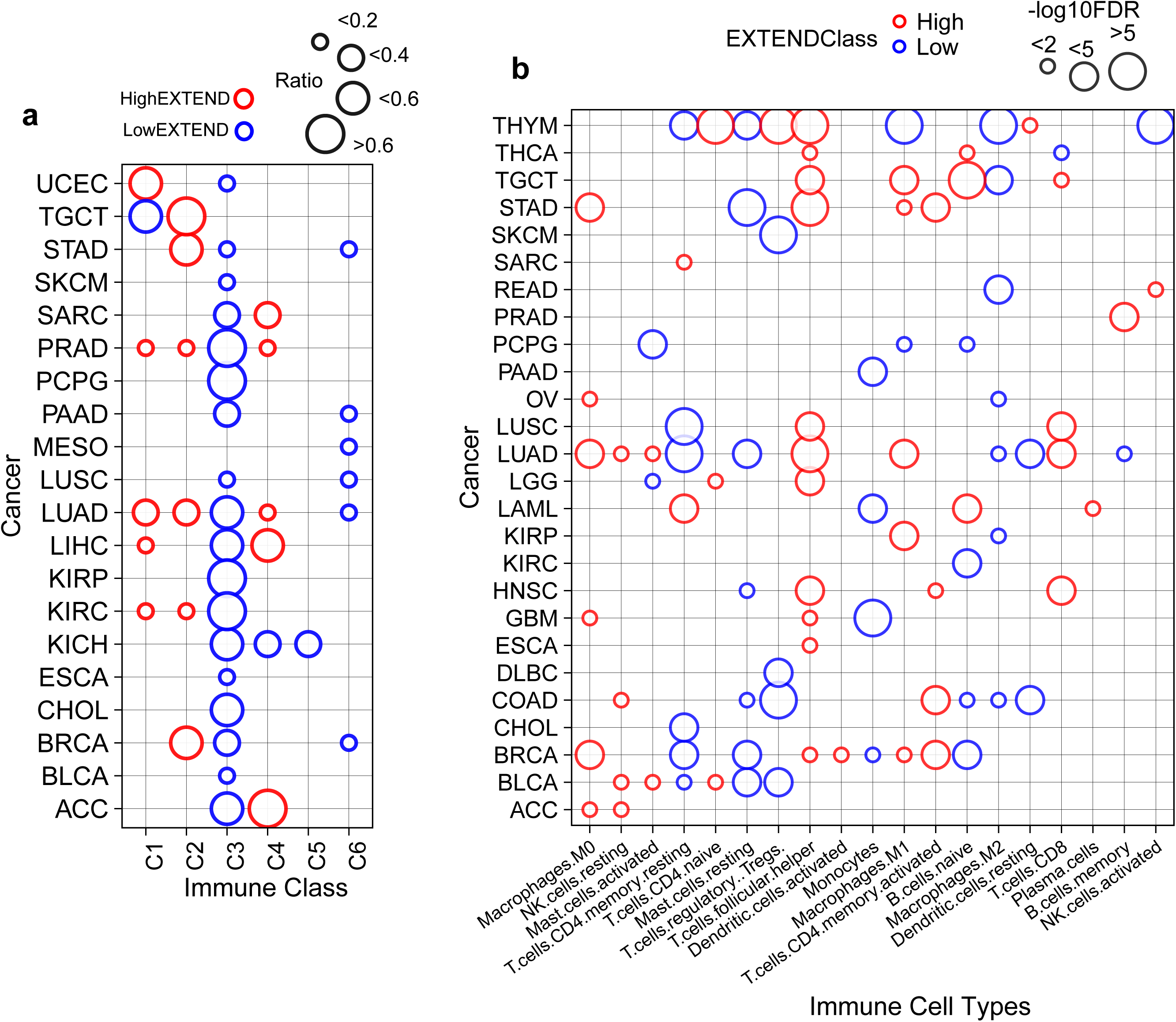
Association of telomerase levels with Immune system. **(a)** Differential fractions of immune classes across low and high telomerase groups in 20 TCGA cancer types. The size of each dot represents the ratio of a respective class **(b)** Immune cell type disparities in low and high telomerase groups. The size of the dots represents -log10 FDR values. The blue color in both (a) and (b) represents low, and the red color represents a high telomerase group.

Next, we studied if these immune class levels could be emulated at the immune cell type level. We used CIBERSORT deconvolution scores for each immune cell type from the TCGA pan-cancer analysis ^95^. We noted that M2 macrophages and resting mast cells are associated with low telomerase class in 7 cancer types. Activated T-cells are majorly associated with high telomerase levels in most cancer histologies, while resting T-cells have low cases of cancer, respectively ^96,97^. In some cancers, macrophages, M0 and M1 were mainly associated with high telomerase class. Monocytes are associated with low telomerase, which is plausible as they are enriched with a high senescence signature (Fig. 7b), corroborating with a previous study where monocytes were shown to be associated with high senescence. ^80^.

Increasing evidence ^98,99^ suggests that the senescence phenotype is majorly facilitated by mitochondrial dysfunction due to reactive oxygen species (ROS). This induces activation of MAPK pathways ^100–102^, which is one of the top enriched pathways for low telomerase activity class across Pancan, as observed in our pathway enrichment analysis mentioned earlier (Supplementary Tables 1 & 2). MAPK activation further results in inflammatory responses along with cell death mechanisms ^101^. Based on our observations, we further investigated how this cycle is associated with telomerase activity on the pan-cancer level. Therefore, we used ROS and MAPK signaling gene signatures from the Molecular Signatures Database (MSigDb) ^64^ and scored them using ssGSEA for 33 TCGA cancer types. By comparing low and high telomerase, MAPK and ROS scores highlighted the significant association of both factors with low telomerase in most cancer histologies (Supplementary Fig. 6a-b, FDR < 0.05). Interestingly, we also observed an intriguing association between ROS and senescence at the pan-cancer level (*Rho* = 0.9, *P* = 9.94-09). This significant correlation was persistent within each cancer type (FDR < 0.05), signifying it to be cancer lineage-independent (Fig. 8a). We also observed the same pattern for ROS and MAPK correlation (FDR <0.05) across each cancer type. Connecting all the pathways leading from the immune inflammation class C3 with low telomerase demarcated strong associations with high senescence and consequently with high ROS and MAPK activities across the pan-cancer TCGA cohort (Fig. 8b).

**Fig. 8:**
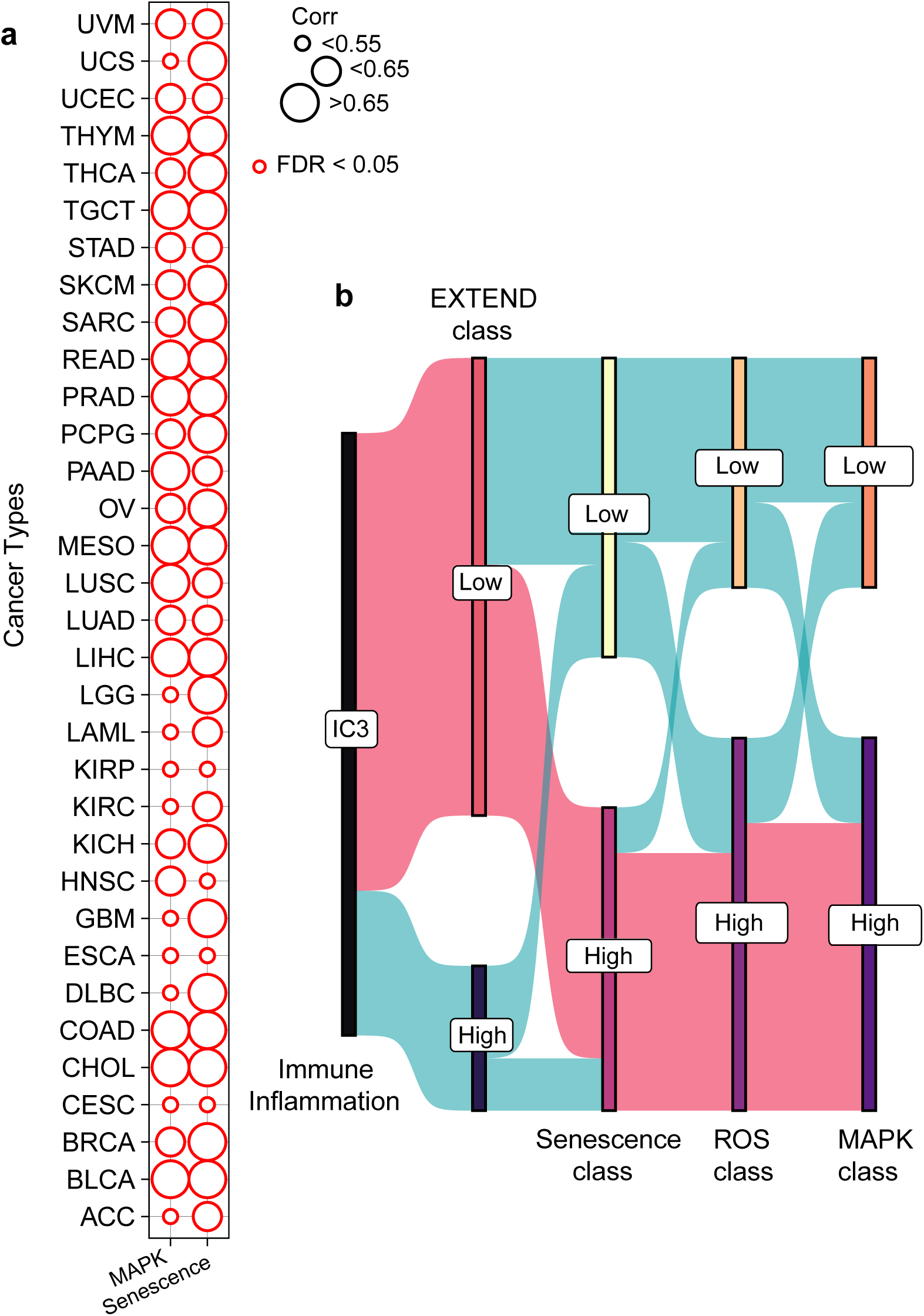
Summary of telomerase associations across multiple cancers. **(a)** Correlation of ROS scores with MAPK and Senescence scores across TCGA Pan cancer. The size of the dots represents correlation coefficients. **(b)** Association summary of telomerase groups with immune, senescence, ROS, and MAPK activities across Pan can.

## Discussion

Telomerase, being a center regulator of all cancer hallmarks ^103^, is studied across different cancers ^19,81,104,105^ but is disregarded for the characterization of its heterogeneous levels across multiple tissue types and how these levels are affected by different genomic features, thereby generating a knowledge gap. We established and reported the method for the quantification of telomerase activities in cancer at bulk and single-cell levels ^29^. To gain further insights into the modulators of telomerase and their impact on its quantitative levels across a cohort of cancers, we proposed a robust classification method.

Using an unsupervised mechanism, we have divided telomerase activity into low and high categories across 33 cancer histologies. In the current study, we systematically investigated the clinical and genomic disparities between low and high telomerase groups. We did not find substantial differences between the two groups in the case of clinical variables, including sex, age, and disease stage, with few exceptions. High telomerase groups generally showed higher tumor mutation burden, gene fusions, and copy number alterations ^48^. Thymoma was an exception to the case. Our previous study ^29^ did not observe any correlation between telomere length ^47^ and telomerase activity. Yet, a comparative analysis of low and high telomerase groups here led to a significant orientation of telomere lengths towards the low telomerase group, with thymoma as an exception. This observation effectively corroborates the association of shorter telomere length with high telomerase and vice versa ^106^.

Division of telomerase activities into low and high classes for >9000 bulk tumors along with single and spatial transcriptomics settings confirmed previous connotations, available in bits and pieces for some cases, and highlighted certain genomic gradients lacking at a genome-wide pan-cancer level. A striking bend of senescence, a measure of cell cycle arrest, towards low telomerase class is explained by their overall low proliferative potential and cancer stemness. Cancer cell lines data and single-cell and spatial transcriptomics analysis further confirmed this trend. The significant association of senescence with low telomerase activity class in older adults in cancer and normal tissue settings highlights the impact of cellular aging on telomerase and senescence in elderly individuals ^73^. This signifies at a pan-cancer level that low telomerase in older adults leads to high senescence and vice versa.

The association of immune inflammation ^94^ with low telomerase class across the majority of cancers indicates its association with the production of reactive oxygen species (ROS) and related mechanisms ^107^. The generation of ROS under the influence of senescence^108,109^ is demonstrated across 33 cancer types. Furthermore, the strong association of MAPK signaling with ROS ^110^ indicates the tight association of these mechanisms. The tight association of immune inflammation class with low telomerase highlights the active mechanisms of senescence, ROS, and MAPK signaling at a pan-cancer level. In summary, our study demonstrates that the unsupervised classification of telomerase levels provides insights into related mechanisms of tumorigenesis. This analysis establishes a histology based significant association of low and high telomerase levels with genomic alterations, senescence, inflammation, and related mechanisms at a pan-cancer level. This has not been defined so far, thus provides a comprehensive view across multiple cancer histologies.

## Methods

### Data Sources

We downloaded the bulk gene expression (in RSEM=RNAseq by Expectation Maximization) and phenotypic data for 33 cancer types from The Cancer Genome Atlas (TCGA) (Synapse ID: syn4874822; https://gdc.cancer.gov/node/905/). These cancer types include ACC, BLCA, BRCA, CESC, CHOL, COAD, DLBC, GBM, HNSC, ESCA, KICH, KIRC, KIRP, LAML, LGG, LIHC, LUAD, LUSC, MESO, OV, PCPG, PRAD, PAAD, SARC, SKCM, STAD, READ, TGCT, THYM, THCA, UCEC, UVM and UCS. Bulk gene expression data for cancer cell lines were downloaded from the Cancer Cell Line Encyclopedia (CCLE; https://portals.broadinstitute.org/ccle, version 02-Jan-2019). Normal tissue bulk gene expression (RNAseq) data was retrieved from Genotype Tissue Expression (GTEx) downloaded as of March 2019 (release 2016-01-15_v7). Bulk gene expression human development data of the liver and heart^89^ were downloaded from Array Express (accession no: E-MTAB-6814). We also used single-cell RNAseq glioblastoma ^83^ and head and neck cancer^84^ data sets in our analysis. Further spatial transcriptomics data for lung ^85^ and breast ^86^ cancers were also used in our study from the mentioned studies. No human subjects were involved in this study.

### Unsupervised Classification of Telomerase Activity Scores

We used EXTEND scores from our previous study ^29^ for all gene expression data sets mentioned in the data sources as a measure of telomerase activity except for two spatial transcriptomics data sets. Telomerase activity scores for spatial data sets were computed using the EXTEND method. We classified EXTEND scores into low and high groups by using an unsupervised K-means ^50^ clustering algorithm. The method was applied per cancer type for all the data sets under study. The K-means algorithm has been implemented using the R platform. The algorithm required EXTEND scores, number of centroids (K), number of iterations and number of random selection of centroids as an input. The centroid (K) referes to a center point for each cluster to be formed. In order to segregate telomerase into low and high levels, K was fixed to 2. In order to attain robust results, the algorithm was iterated 1000 times with 50 randomly selected centroids(K). The final classification of samples/cells into low and high telomerase groups was based on the consensus of 50,000 iterations.

### Clinical and Demographic Variables Comparisons

We associated high and low telomerase groups with various factors to identify potential regulators of telomerase. These regulators include clinical and demographic features such as gender, age, disease stage, and overall survival for TCGA data. Telomerase groups were compared with age using two classifications, first by dividing the samples into young adults (18-50 years) and older adults (>50 years) to generate a balanced set for comparison purposes across TCGA. Secondly, a more comprehensive analysis of age with telomerase was performed on TCGA and GTEX data to divide the samples into younger adults (AYA) and older adults (OA) based on a recent study ^111^, i.e., between 15-39 are marked as AYA and > 39 were marked as OA. The samples with no age information, pediatric samples with very few cases, and cancer types with less than 15 cases were removed from comparisons. We compared these features using Fisher’s exact test followed by multiple hypothesis testing using the FDR method.

### Genome instability comparisons

We also compared low and high telomerase classes across each cancer type in TCGA based on various genomic features. These include telomere lengths (TL) retrieved from a previous study^47^. We compared the tumor to the normal ratio of telomere lengths (TL ratio) as a measure of comparison. We further compared tumor mutation burden (TMB), copy number altered fractions (CNAs), whole genome doubling (WGD), and gene fusion-based genomic abnormalities among low and high telomerase groups across all cancer types. The data for TMB, CNVs, and WGD was retrieved from gdc.cancer.gov (https://gdc.cancer.gov/about-data/publications/panimmune). TCGA gene fusions and druggable fusions data were downloaded from the TCGA resource study ^66^. Differential analysis for the genomic aberrations was conducted using a two-sided t-test and fisher-exact test where applicable, followed by multiple hypothesis testing using the FDR method.

### Senescence scores comparison

We calculated senescence scores for bulk RNAseq data sets, including TCGA, GTEX, CCLE, and human development data, using ssGSEA implemented in R package “GSVA” ^112^. The senescence scores were calculated using a gene set termed as SenMayo comprising 125 genes from one of the recent studies ^80^. Single-cell GBM and HNSC data sets were processed using the Seurat ^113^ toolkit function “Addmodule score” to calculate senescence scores. Spatial transcriptomics-filtered data sets for both lung and breast cancer were used for score calculation using the AUCell ^114^ method. Differential analysis for senescence scores between low and high telomerase groups for all the data sets was conducted using a two-sided t-test. The *P* values were subjected to multiple testing using FDR correction. Further correlation of telomerase activity and senescence scores was calculated using spearman-based method for all the data sets.

Senescence scores for low and high telomerase groups were compared between older adults for 33 cancer types in the TCGA data. The cancer types with matched normal tissues in GTEX data with a significant sample size >50 were utilized for age comparisons using a t-test. Due to the small sample size of human development data, a Spearman-based association was utilized to identify the trend between senescence and telomerase activities.

### Immune Cell Types and their association

We used immune classes identified based on immune cell properties of cancer tissues by TCGA resource studies ^94^. For TCGA data, we calculated the association of high and low telomerase groups with all immune classes. Furthermore, we used immune cell types predicted by ESTIMATE for TCGA data and identified differential patterns for high and low telomerase cases. Immune classes were further associated with multiple scores, including telomerase, senescence, MAPK, and ROS, to highlight the significant association among these pathways. The gene sets for MAPK and ROS as were retrieved from MSigDB and were scored using ssGSEA. The score comparisons were performed using a t-test followed by multiple tests using the FDR method.

## Declarations

### Ethical Approval

This declaration is not applicable

### Author contributions

N.N. and M.H.K. conceptualized and designed the research; N.N collected and analyzed the data; N.N. and M.H.K. wrote the manuscript.

## Data and Materials Availability

All the data used in this project is from publicly available resources and is mentioned in the Data Sources in the Methods Section. Processed data used to generate the main and supplementary figures will be provided as Source Data File upon acceptance. Scripts used to generate the figures are available at the GitHub link https://github.com/NNoureen/Categorization

---

**Supplementary Fig. 1.**
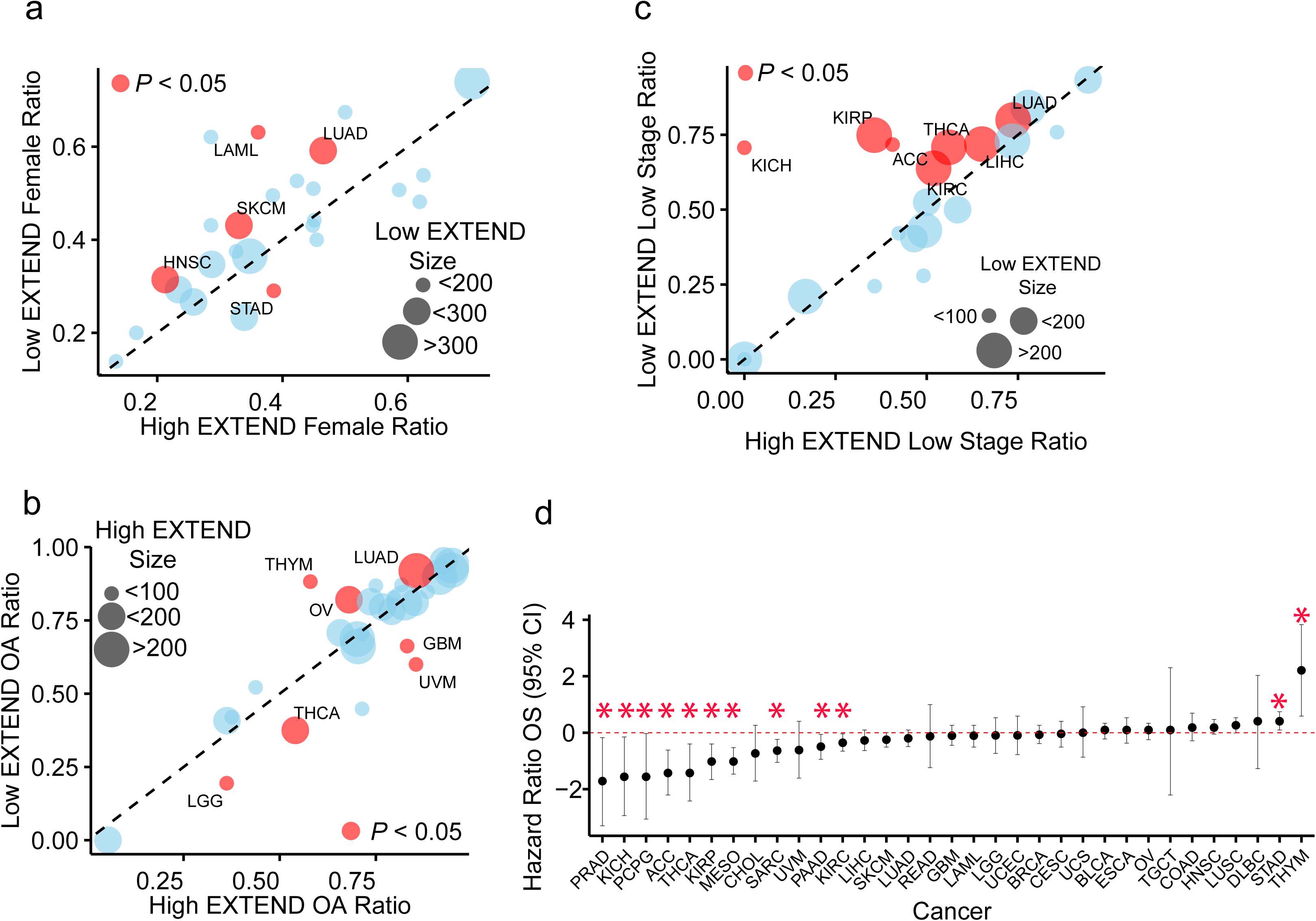
Clinical disparities between low and high telomerase groups. **(a)** Female fraction **(b)** Older adults fraction **(c)** and Low Stage fraction comparisons between high and low telomerase groups. Cancer types are labelled for significant cases (P < 0.05; Fisher exact test). The data size reflects low and high extend classes size for representative cases. **(d)** Hazard ratio plot (y-axis = log scale) based on low and high extend groups for 31 cancer types. Significant cases(P< 0.05) are marked with asterisk.

**Supplementary Fig. 2.**
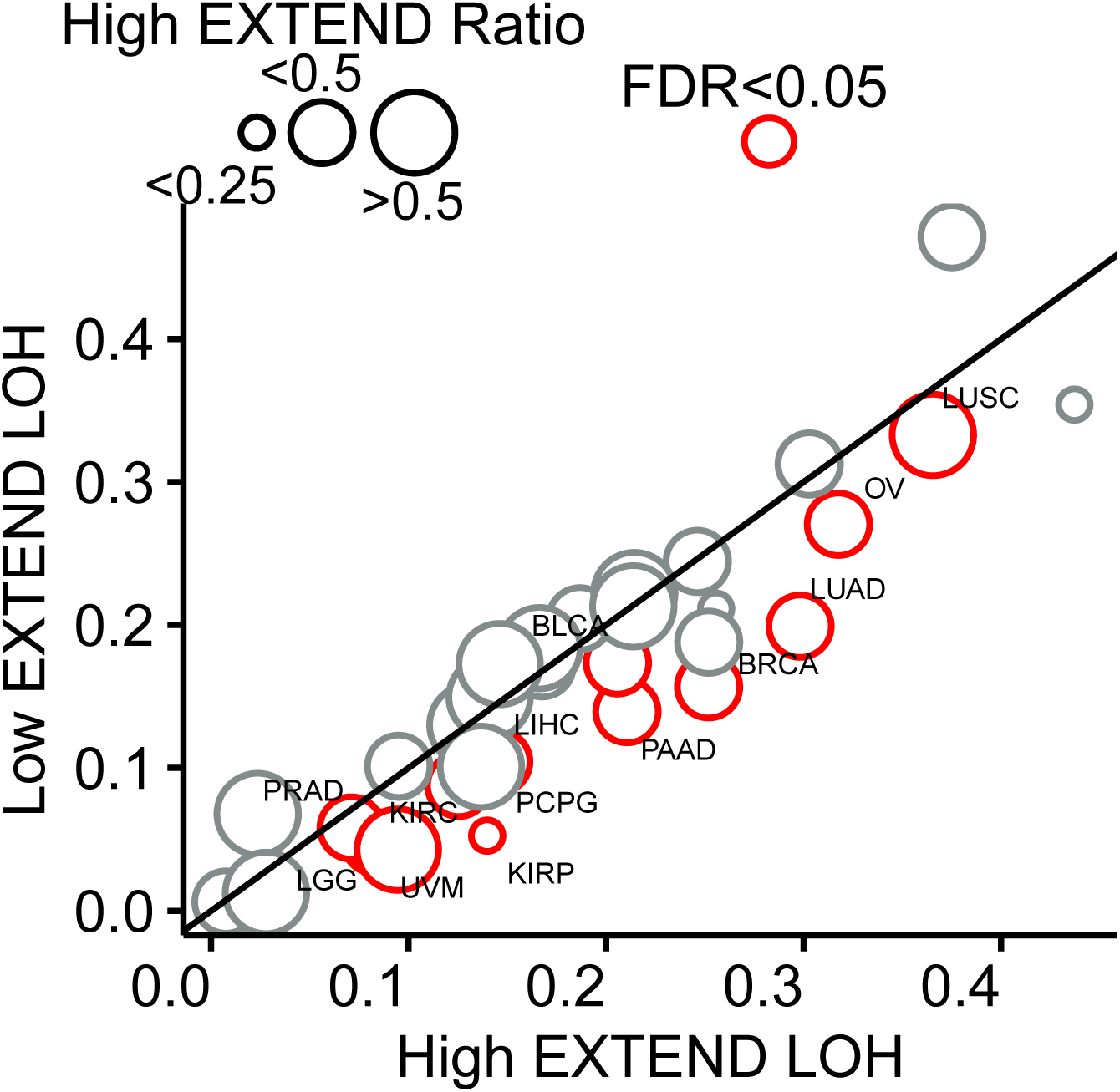
Genomic disparities for low and high telomerase across Pancan. Differential patterns of loss of heterozygosity (LOH), for low and high telomerase groups across Pan-cancer. Significance is shown as FDR < 0.05 (red color) and size of the data shows ratio of high telomerase group for each cancer type.

**Supplementary Fig. 3.**
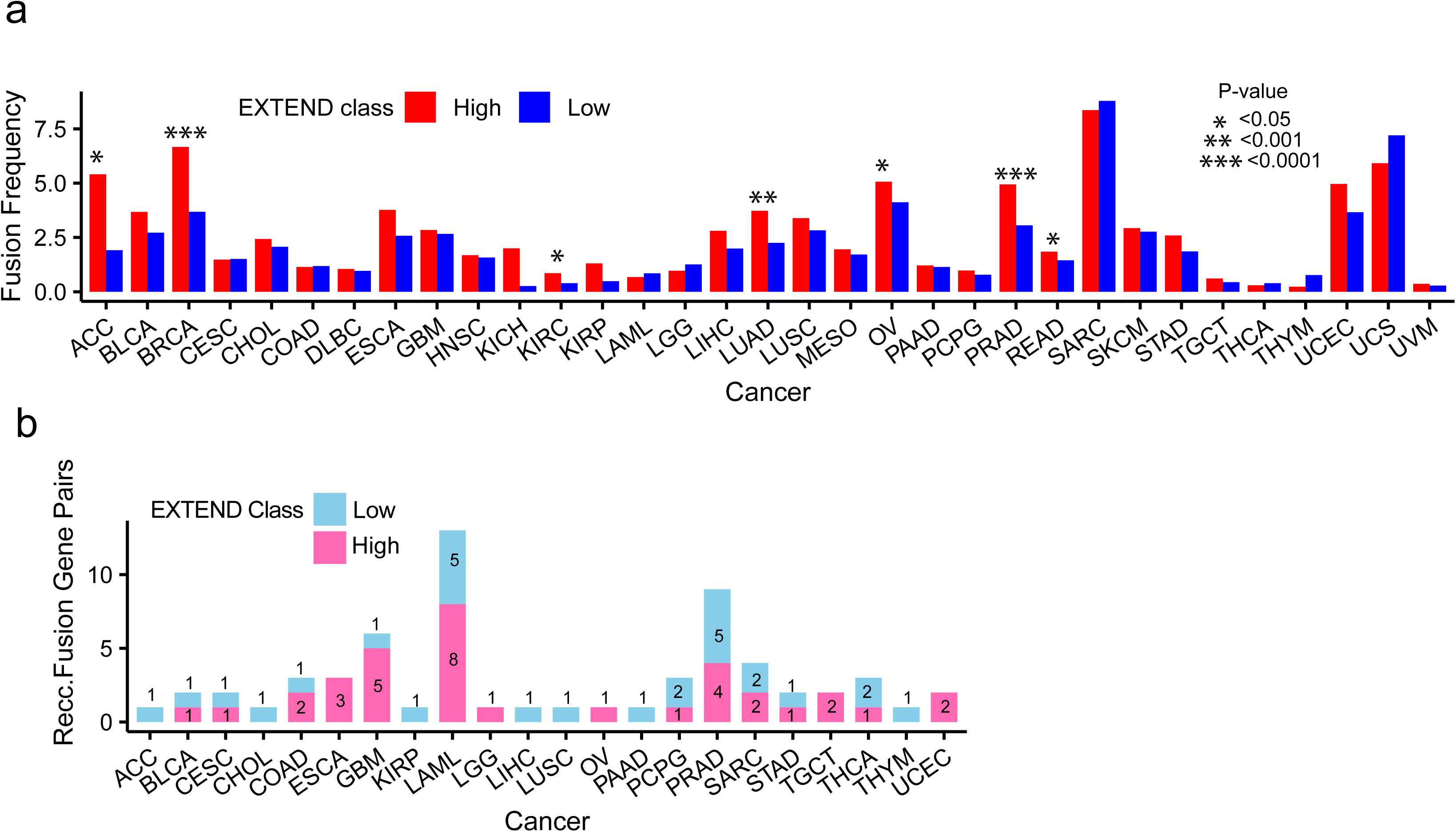
Gene fusion frequencies for low and high telomerase across Pancan. Differential patterns of **(a)** Overall gene fusion frequencies distributed across low (blue) and high (red) telomerase groups for Pancan. **(b)** Recurrent fusion frequencies for each gene-pair for low (blue) and high (pink) telomerase group across pan-cancer cohort. The numbers on each bar reflect the total number of gene-pairs for each case. Fusions with greater than 1 percent of cases per cancer type are included in analysis.

**Supplementary Fig. 4.**
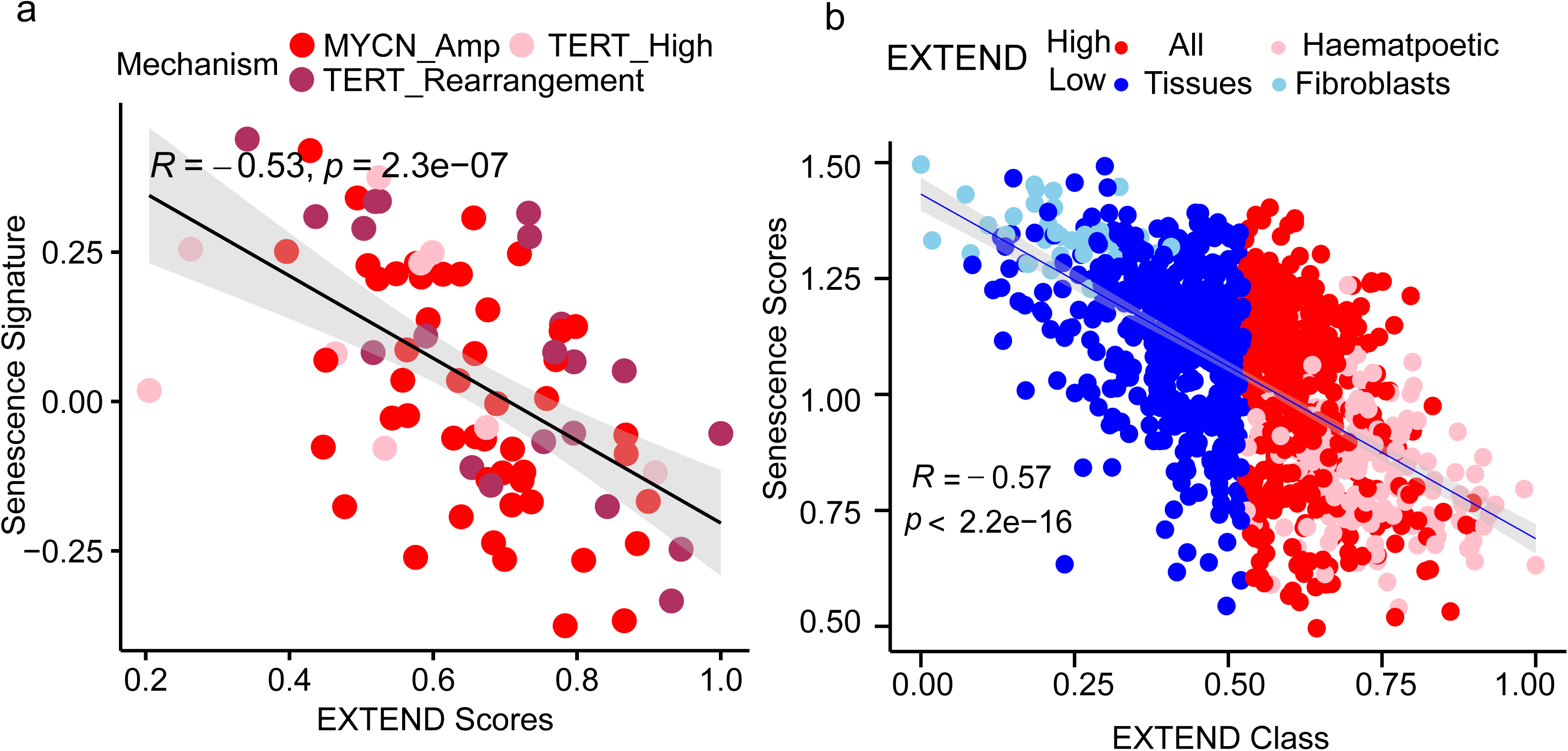
Association of telomerase with senescence Correlation (Spearman) of telomerase (EXTEND) and senescence scores in **(a)** Neuroblastoma data (Ackerman *et al*., 2018) **(b)** CCLE data; top low (light blue = fibroblasts) and high (pink = hematopoietic) scoring cell lines are marked separately, remaining are highlighted as low (blue) and high (red)

**Supplementary Fig. 5.**
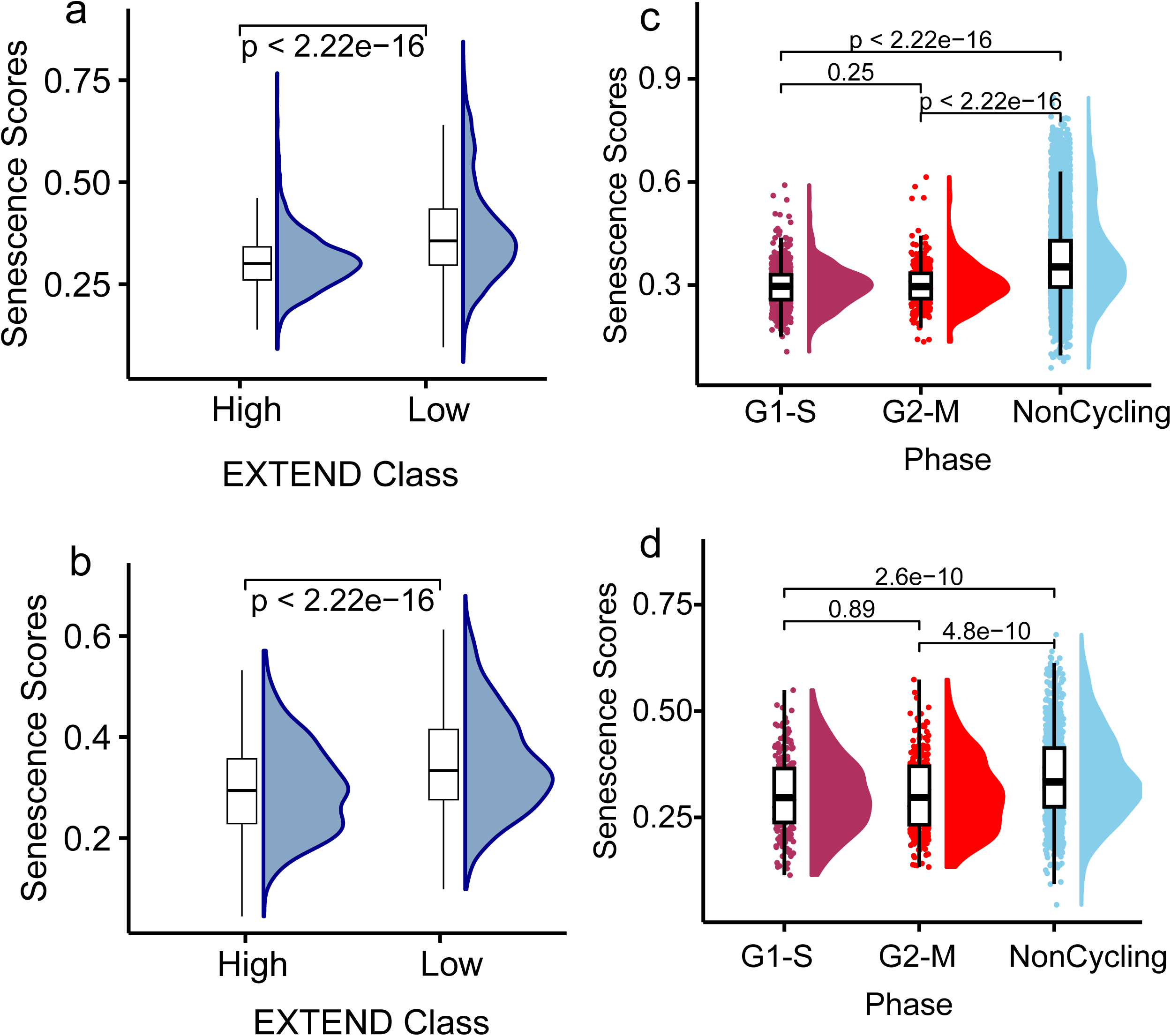
Differential patterns of senescence in low and high telomerase groups for single cell data. Senescence scores differences between low and high telomerase groups in **(a)**Single cell GBM data **(b)** Single cell HNSC data. Senescence scores patterns distributed across cell cycle phases for (c) Single cell GBM data **(d)** Single cell HNSC data. *P* values are calculated using Student’s T-test.

**Supplementary Fig. 6.**
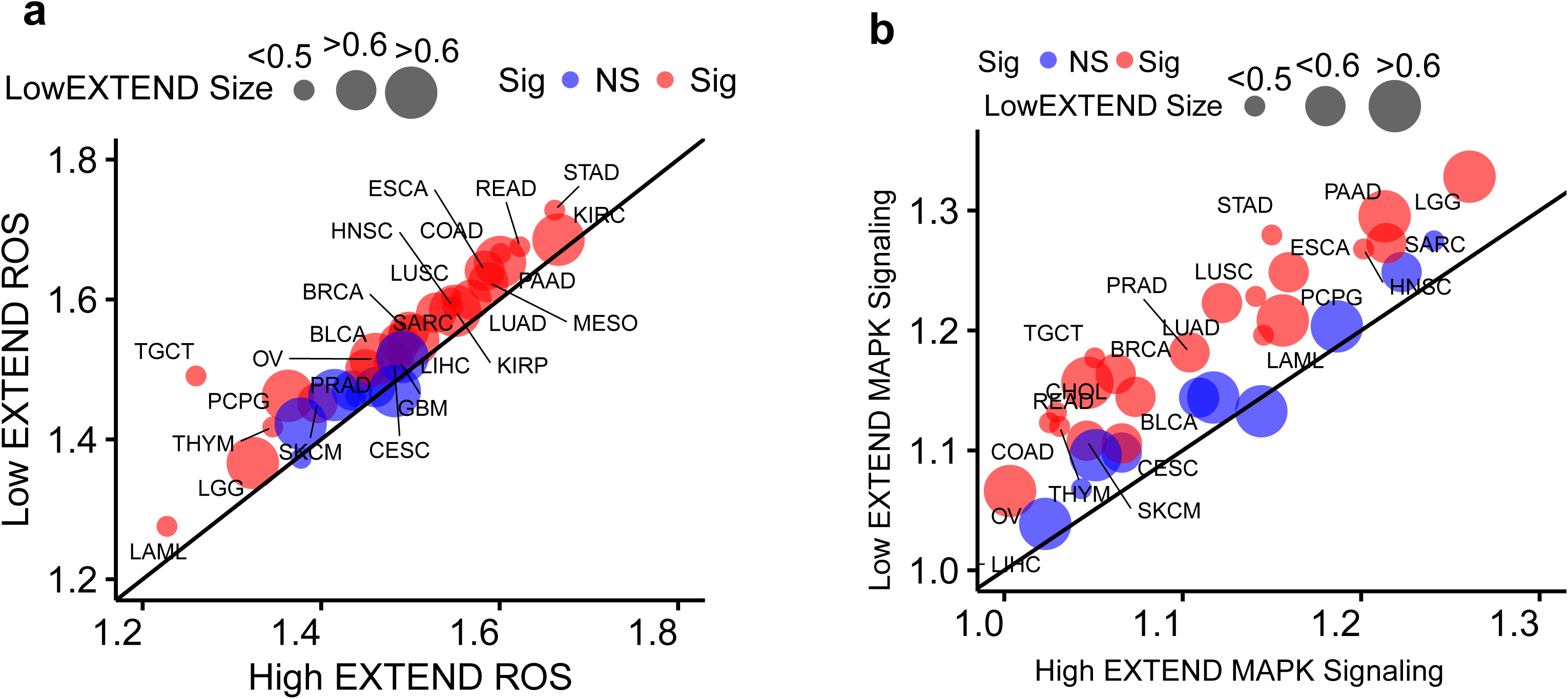
ROS and MAPK Signaling across Telomerase Groups. Differential patterns of low and high telomerase classes across TCGA pan-cancer data for **(a)** ROS scores **(b)** MAPK signaling scores. Significance level (red) shown as FDR < 0.05. Size of the data represents low EXTEND scores class size.

**Supplementary Table 1:** High EXTEND class gene set enrichment analysis for Pan cancer.

| Gene Set Name | Pvalue | FDR | EXTEND Class |
| --- | --- | --- | --- |
| KORKOLA_CORRELATED_WITH_POU5F1 | 4.27E-18 | 9.96E-14 | High |
| CONRAD_STEM_CELL | 3.20E-17 | 3.72E-13 | High |
| GOBP_DEVELOPMENTAL_PROCESS_INVOLVED_IN_REPRODUCTION | 2.07E-14 | 1.61E-10 | High |
| GOBP_GAMETE_GENERATION | 1.08E-13 | 5.26E-10 | High |
| GOCC_PROTEIN_DNA_COMPLEX | 1.13E-13 | 5.26E-10 | High |
| GOBP_SEXUAL_REPRODUCTION | 9.67E-13 | 3.32E-09 | High |
| BENPORATH_ES_2 | 9.99E-13 | 3.32E-09 | High |
| GOBP_MULTICELLULAR_ORGANISM_REPRODUCTION | 1.91E-12 | 5.57E-09 | High |
| GOBP_REPRODUCTION | 3.81E-12 | 9.86E-09 | High |
| GOCC_CHROMOSOME | 4.85E-12 | 1.13E-08 | High |
| KORKOLA_EMBRYONAL_CARCINOMA | 8.71E-12 | 1.84E-08 | High |
| GOBP_MALE_GAMETE_GENERATION | 1.32E-11 | 2.56E-08 | High |
| REACTOME_TRANSCRIPTIONAL_REGULATION_OF_PLURIPOTENCY | 2.34E-11 | 4.02E-08 | High |
| CONRAD_GERMLINE_STEM_CELL | 2.42E-11 | 4.02E-08 | High |
| GOBP_GERM_CELL_DEVELOPMENT | 6.87E-11 | 1.07E-07 | High |
| GOBP_CELLULAR_PROCESS_INVOLVED_IN_REPRODUCTION | 7.37E-11 | 1.07E-07 | High |
| BENPORATH_ES_1 | 1.22E-10 | 1.67E-07 | High |
| WP_CARDIAC_PROGENITOR_DIFFERENTIATION | 7.06E-10 | 9.13E-07 | High |
| GOMF_SEQUENCE_SPECIFIC_DNA_BINDING | 1.18E-09 | 1.44E-06 | High |
| WP_PRE_IMPLANTATION_EMBRYO | 1.23E-09 | 1.44E-06 | High |
| GOBP_GONADAL_MESODERM_DEVELOPMENT | 1.35E-09 | 1.50E-06 | High |
| PASINI_SUZ12_TARGETS_UP | 1.75E-09 | 1.85E-06 | High |
| MIKKELSEN_PLURIPOTENT_STATE_UP | 3.54E-09 | 3.58E-06 | High |
| WP_CELL_LINEAGE_MAP_FOR_NEURONAL_DIFFERENTIATION | 5.51E-09 | 5.35E-06 | High |
| GOBP_MESODERM_DEVELOPMENT | 6.79E-09 | 6.32E-06 | High |
| GOBP_PROTEIN_DNA_COMPLEX_ORGANIZATION | 8.20E-09 | 7.34E-06 | High |
| GOMF_CHROMATIN_BINDING | 1.36E-08 | 1.18E-05 | High |
| GOMF_HISTONE_BINDING | 2.38E-08 | 1.98E-05 | High |
| REACTOME_GERM_LAYER_FORMATION_AT_GASTRULATION | 2.53E-08 | 2.03E-05 | High |
| REACTOME_DEVELOPMENTAL_BIOLOGY | 7.10E-08 | 5.51E-05 | High |
| MIKKELSEN_ES_ICP_WITH_H3K4ME3 | 1.05E-07 | 7.85E-05 | High |
| REACTOME_MATERNAL_TO_ZYGOTIC_TRANSITION_MZT | 1.49E-07 | 1.09E-04 | High |
| GOBP_NUCLEOSOME_ORGANIZATION | 2.33E-07 | 1.65E-04 | High |
| GOBP_CHROMATIN_REMODELING | 2.94E-07 | 2.01E-04 | High |
| GOBP_REGULATORY_NCRNA_PROCESSING | 6.24E-07 | 4.15E-04 | High |
| MATZUK_SPERMATID_DIFFERENTIATION | 6.83E-07 | 4.42E-04 | High |
| GOBP_CELL_FATE_COMMITMENT | 1.08E-06 | 6.77E-04 | High |
| HADDAD_B_LYMPHOCYTE_PROGENITOR | 1.18E-06 | 7.23E-04 | High |
| GOMF_DNA_BINDING_TRANSCRIPTION_FACTOR_ACTIVITY | 2.55E-06 | 1.38E-03 | High |
| MALTA_CURATED_STEMNESS_MARKERS | 7.90E-06 | 3.54E-03 | High |
| BHATTACHARYA_EMBRYONIC_STEM_CELL | 1.87E-05 | 7.01E-03 | High |
| GOBP_MAINTENANCE_OF_CELL_NUMBER | 2.64E-05 | 9.18E-03 | High |

**Supplementary Table 2:**
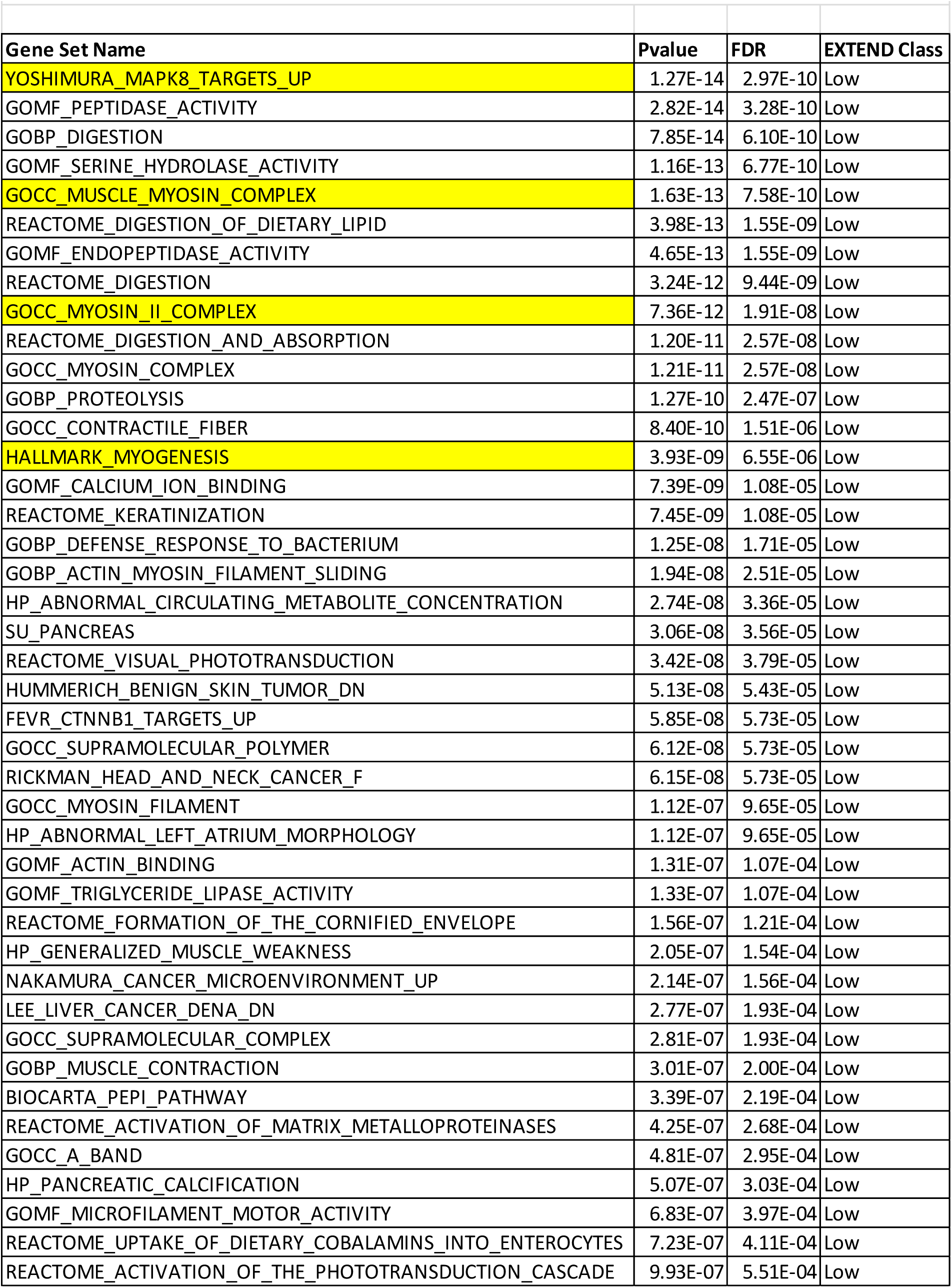
Low EXTEND class gene set enrichment analysis for Pan cancer.

